# Mixed-stock analysis in the age of genomics: Rapture genotyping enables evaluation of stock-specific exploitation in a freshwater fish population with weak genetic structure

**DOI:** 10.1101/2020.11.10.376350

**Authors:** Peter T. Euclide, Tom MacDougall, Jason M. Robinson, Matthew D. Faust, Chris C. Wilson, Kuan-Yu Chen, Elizabeth A. Marschall, Wesley Larson, Stuart A. Ludsin

## Abstract

Mixed-stock analyses using genetic markers have informed fisheries management in cases where strong genetic differentiation occurs among local spawning populations, yet many fisheries are supported by multiple spawning stocks that are weakly differentiated. Freshwater fisheries exemplify this problem, with many harvested populations supported by multiple stocks of young evolutionary age that are isolated across small spatial scales. As a result, attempts to conduct genetic mixed-stock analyses of inland fisheries have often been unsuccessful. Advances in genomic sequencing now offer the ability to discriminate among populations with weak population structure by providing the necessary resolution to conduct mixed-stock assignment among previously indistinguishable stocks. We demonstrate the use of genomic data to conduct a mixed-stock analysis of Lake Erie’s commercial and recreational walleye (*Sander vitreus*) fisheries and estimate the relative harvest of weakly differentiated stocks (pairwise *F*_ST_ < 0.01). We used RAD-capture (Rapture) to sequence and genotype individuals at 12,081 loci that had been previously determined to be capable of discriminating between western and eastern basin stocks with 95% reassignment accuracy, which was not possible in the past with microsatellite markers. Genetic assignment of 1,075 fish harvested from recreational and commercial fisheries in the eastern basin indicated that western basin stocks constituted the majority of individuals harvested during peak walleye fishing season (July – September). Composition of harvest changed seasonally, with eastern basin fish comprising much of the early season harvest (May – June). Clear spatial structure in harvest composition existed; more easterly sites contained more individuals of east basin origin than did westerly sites. Our study provides important stock contribution estimates for Lake Erie fishery management and demonstrates the power of genomic data to facilitate mixed-stock analysis in exploited fish populations with weak population structure or limited existing genetic resources.

## Introduction

The sustainability of many populations depends on multiple, but often cryptic, breeding groups connected by shared habitat or reproductive behaviors (Hilborn et al. 2003; Cowen et al. 2007; Alves et al. 2010). Such complexity helps to improve population stability and resilience (Hilbron et al. 2003; Schindler et al. 2010; 2015), but it also complicates conservation and management by increasing the number of areas where conservation and regulatory actions could occur (Cooke et al. 2016). In some cases, populations have been sustainably managed by treating discrete local spawning stocks as parts of an overall portfolio of population diversity (Schindler et al. 2015; Waples et al. 2020). Although this idea has primarily been applied to marine fisheries, it has also been suggested to be relevant to freshwater fisheries (DuFour et al. 2015). However, fisheries management often presumes that assessment information represents a single stock as opposed to multiple stocks, which can lead to unintended overexploitation of local stocks and inappropriate management advice (Stephenson 1999; Li et al. 2015). For example, the collapse of Atlantic cod *Gadus morhua* populations is thought to have been caused in part by a failure to account for multiple stocks in the assessment and management of this fishery (Hutchinson 2008). An ability to accurately discriminate and identify population components (such as local spawning stocks) is integral to the conservation and management of populations that fit the portfolio theory model of ecology and evolution because it provides a means of accounting for variance in stock-specific productivity (Figge 2004; Sethi 2010). However, achieving accurate discrimination among population components can be difficult, especially for freshwater species in large lakes that often experience high gene flow similar to marine populations, but at reduced spatial scales (Martinez et al. 2018) or are of young evolutionary age. Thus, the need exists for methods that can deal with the resultant weak population structure.

For decades, molecular markers have been used to resolve stock-specific contributions to mixed-stock fisheries in marine systems (Seeb and Crane 1999; Milner et al. 2008; Bernatchez et al. 2017; Waples et al. 2020). For example, mixed-stock assessments have become central to the management of Pacific salmonines (Shaklee et al. 1999) such that the contributions of hundreds of salmon spawning populations are evaluated annually using molecular markers (Dann et al. 2013; Beacham et al. 2020). Similar practices have become essential to Atlantic Cod *Gadus morhua* management by improving understanding of seasonal spawning dynamics (Dean et al. 2019) and limiting overharvest of less productive stocks (Dahle et al. 2018). Freshwater fisheries could benefit from similar portfolio-based approaches to management as they are often under high fishing pressure (DuFour et al. 2015; Fluet-Chouinard et al. 2018; Embke et al. *in press*) and experience more environmental stochasticity than marine systems (Strayer and Dudgeon 2010). However, application of mixed-stock assessments use of similar techniques has been limited, partially owing to difficulties in developing molecular resources capable of accurately discriminating among stocks (e.g., Chen et al. 2020, but see Andvik et al. 2016).

Advancing stock identification in freshwater systems is important because the same portfolio theory strategies used in marine systems have been suggested to improve population stability of exploited freshwater species that display complex portfolios of diversity (DuFour et al. 2015). Because the habitat of many freshwater species shifted southward and was reduced during the last Ice Age, the recolonized populations remain relatively young (4,000 – 12,000 years old) and lack substantial genetic differences among spawning stocks (Bernatchez & Wilson, 1998). Additionally, freshwater populations often contain spawning stocks in relatively close proximity to each other and admixture among stocks can complicate stock reassignment (Waples and Gaggiotti 2006; Martinez et al. 2018). Therefore, traditional genetic approaches such as microsatellites often lack the statistical power to consistently discriminate among spawning stocks (Johnson et al. 2004). Instead, the development of high-resolution genomic marker panels that can discriminate among local spawning stocks could make mixed-stock management accessible for fisheries managers tasked with managing such species.

Developing high-resolution marker panels can require a large amount of time and analytical experience, which has limited the use of such panels to high-value marine fisheries, such as salmonids and cod. However, the cost of conducting genomic studies has decreased substantially (Meek and Larson 2019) and molecular and statistical advancements have made it possible to untangle complex patterns of population structure and conduct stock identification of mixed-stock harvests (Layton et al. 2020; Carrier et al. 2020). For example, advances in genotyping-by-sequencing technology, such as RAD-capture (Rapture), have made it possible to affordably and consistently genotype thousands of genetic markers to discriminate among finely structured populations (Ali et al. 2016; Sard et al. 2020). Rapture is a sequence capture based approach that uses biotinylated RNA capture baits to target loci identified using reducedLrepresentation approaches, such as restriction site associated DNA sequencing (RAD-Seq). Additionally, new bioinformatic approaches have led to increased discriminatory power among populations with weak genetic structure. For example, the identification and use of microhaplotypes, which are polyallelic haplotypes of two or more SNPs that are in high linkage disequilibrium and can be sequenced together within the same 150-300 bp region (McKinney et al. 2017a; Baetscher et al. 2018). Advances in computing power and statistics have also resulted in improved reassignment accuracy through resampling cross-validation and machine-learning algorithms (Anderson et al. 2008; Chen et al. 2018). As a result, mixed-stock assignment using genomic data offers the potential to benefit management of fisheries supported by stocks that are difficult to distinguish and that are vulnerable to differentiated overexploitation by commercial, recreational, and subsistence fishing (Post et al. 2002; Fluet-Chouinard et al. 2018).

Fish populations supported by multiple stocks are common in the Laurentian Great Lakes, which have a long history of commercial, recreational, and subsistence fishing (Regier and Hartman 1973; Lynch et al. 2016). Walleye (*Sander vitreus*) are one of the most ecologically and economically important species in the Great Lakes (Hatch et al. 1987; Ludsin et al. 2014) and has been the focus of many previous stock discrimination research efforts (Johnson et al. 2004; Stepien et al. 2015; Chen et al. 2017). Lake Erie’s walleye population is the largest of the five Laurentian Great Lakes and is supported by multiple discrete stocks that spawn throughout the lake (Stepien and Faber 1998; Zhao et al. 2011; Stepien et al. 2012). Most of the lake’s walleye production occurs in the western basin where tributary and open-lake spawning stocks of varying productivity exist (DuFour et al. 2015). Individuals from these stocks move throughout Lake Erie during non-spawning periods in search of preferred habitat (Kersher et al. 1999; Raby et al 2018), where they intermix with the smaller spawning stocks from the central and eastern basins (Zhao et al. 2011; Vandergoot and Brenden 2014; Matley et al. 2020). Migration of western basin stocks into Lake Erie’s eastern basin is likely to have a disproportionate influence on local commercial and recreational fisheries because of presumed differences in population productivity and abundance (Zhao et al. 2011). However, the degree to which western versus eastern stocks are harvested by recreational and commercial fisheries in a given year or season is unknown. Further, efforts to discriminate among western and eastern stocks to facilitate mixed-stock assessments using biological markers have been largely unsuccessful (Riley and Carline 1982; Hedges 2002; Johnson et al. 2004; Chen et al. 2017). These needs, in turn, limit management in Lake Erie (MF, JR, TM *personal communication*).

Genomic tools represent an affordable solution for understanding stock-specific dynamics with a recent study using thousands of single nucleotide polymorphisms (SNPs) genotyped with RAD-seq suggesting that high-accuracy reassignment of walleye to basin of origin is possible using genomic tools (Chen et al. 2020). Here, we developed a Rapture panel containing thousands of genetic markers to conduct a mixed-stock analysis of walleye harvest in Lake Erie’s eastern basin. Specifically, our objectives were to 1) quantify the relative contributions of western and eastern basin walleye populations to commercial and recreational harvest within the eastern basin, and 2) determine if the contributions of western and eastern basin populations varied spatio-temporally within the eastern basin. We hypothesized that walleye from the more productive western basin spawning stocks would comprise the majority of harvest for both types of fisheries, but that harvest composition would vary spatially and temporally based on annual seasonal migrations (Kershner et al. 1999; Raby et al. 2018). In testing this hypothesis, we ultimately demonstrate the ability of genomic data to resolve genetic stock structure and facilitate mixed-stock analysis in systems with weak population structure, to the benefit of fisheries management.

## Materials and Methods

### Study system

Lake Erie, which has a surface area of ~ 26,000 km^2^, is composed of three distinct basins. The western basin is shallowest with most areas ranging 3 – 7 m in depth. The central basin is deeper (15 – 18 m deep), with the eastern basin being the deepest (15 – 25 m) and also containing the deepest point in the lake (65 m; Holcombe et al. 2003). Lake Erie supports the largest commercial and recreational walleye fisheries in the world, which are collectively valued at more than 1 billion dollars annually (Roseman et al. 2014). These fisheries are managed via inter-jurisdictional fisheries management with an annual total allowable catch that is distributed across the lake without consideration of relative stock contributions (Kayle et al. 2015; Fig. 1).

**Fig. 1:**
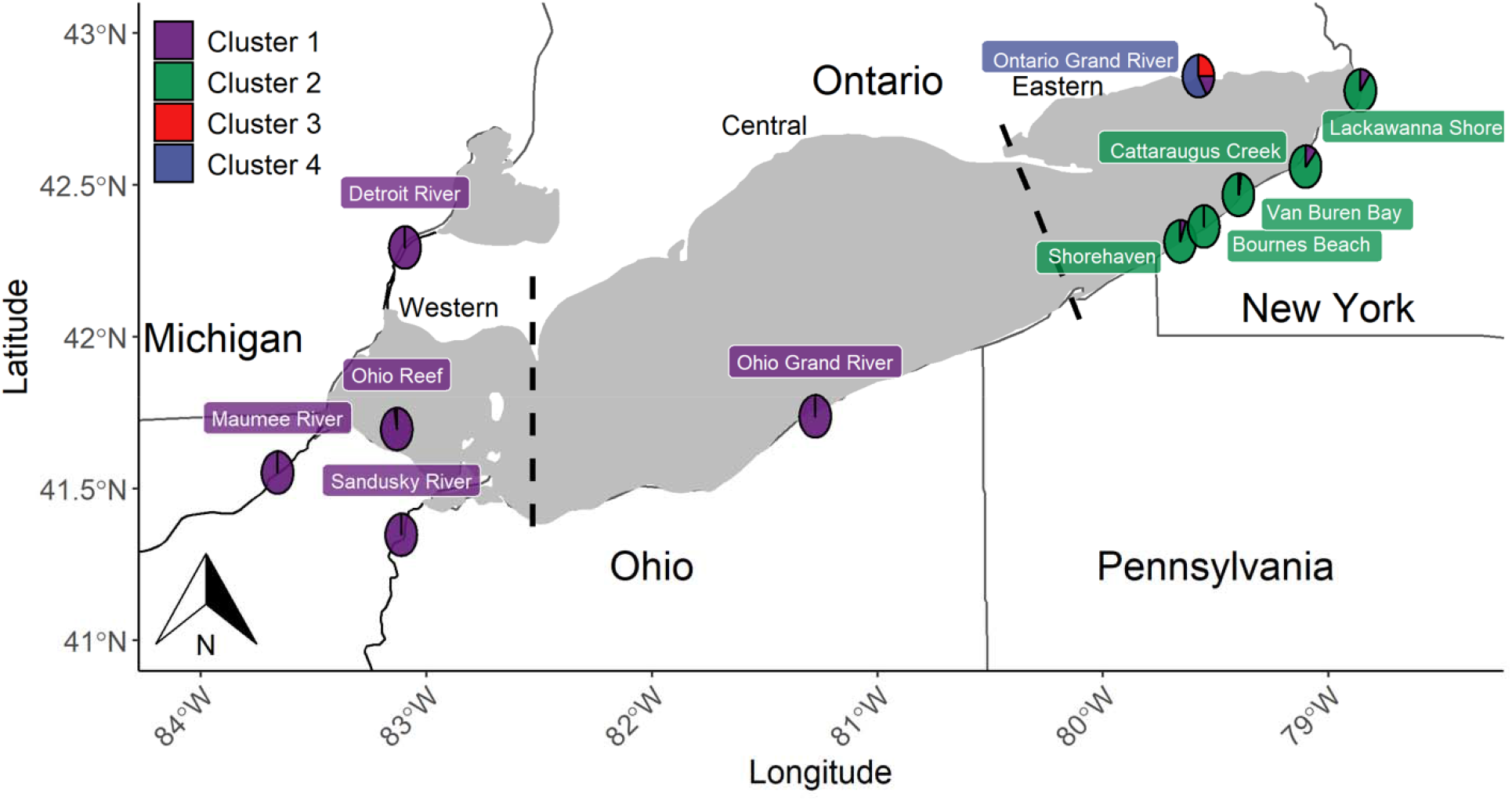
Baseline sample locations and discriminant analysis of principal components (DAPC) results for walleye clustered into the four most parsimonious groups based on Bayesian Information Criterion. Pie charts display the posterior probability of each DAPC cluster proportion averaged across all individuals at the site. The background color of the site label indicates the final reporting group into which the stock was grouped for assignment tests. Lake basins are delineated by dotted lines and labeled on the northern shore of the lake.

In the western basin, four large stocks exist. The largest stocks spawns on open-lake reefs, whereas the others spawn in three large tributaries (Maumee, Sandusky, and Detroit Rivers) that drain into the western basin (DuFour et al. 2015; Fig. 1). Spawning is known to occur in the central basin on nearshore reefs and in the Grand River, Ohio (Stepien et al. 2018). Although the contributions of the resulting offspring to the lake-wide fishery remains unknown for these stocks, it is suspected to be much less than that from the western basin stocks. More spawning stocks exist in the eastern basin, where small spawning aggregations have been reported in small New York tributaries and nearshore reefs dispersed along the south-eastern shore of the east basin, as well as in Ontario’s Grand River on the northern shore of the eastern basin (Zhao et al. 2011; Fig. 1).

### Rapture panel development

We developed a Rapture bait panel (Ali et al. 2016) to reduce the cost and increase the power and consistency of genetic stock identification. Loci for our Rapture panel were identified by conducting a preliminary RAD-sequencing on 96 walleye (Table 1). These individuals were sampled on known spawning grounds during the peak spawning season and were and therefore presumed to have originated from that site (Chen et al. 2020). All of these fish were collected by Lake Erie management agencies, including: Ohio Department of Natural Resources, New York Department of Environmental Conservation, and Ontario Ministry of Natural Resources and Forestry. Fin clips were preserved in 95% ethanol until DNA extraction using Qiagen DNEasy 96 kits.

**Table 1:**
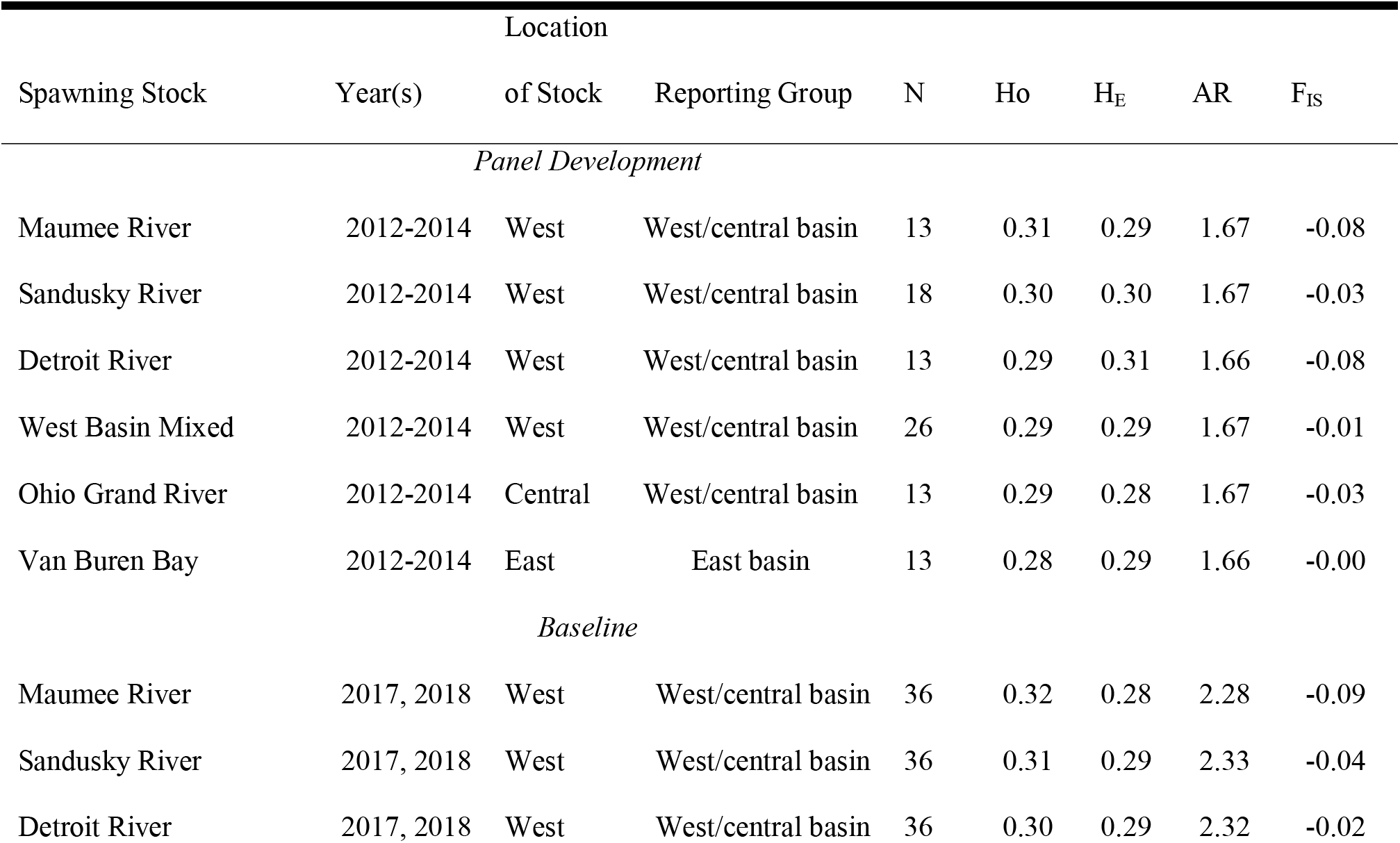
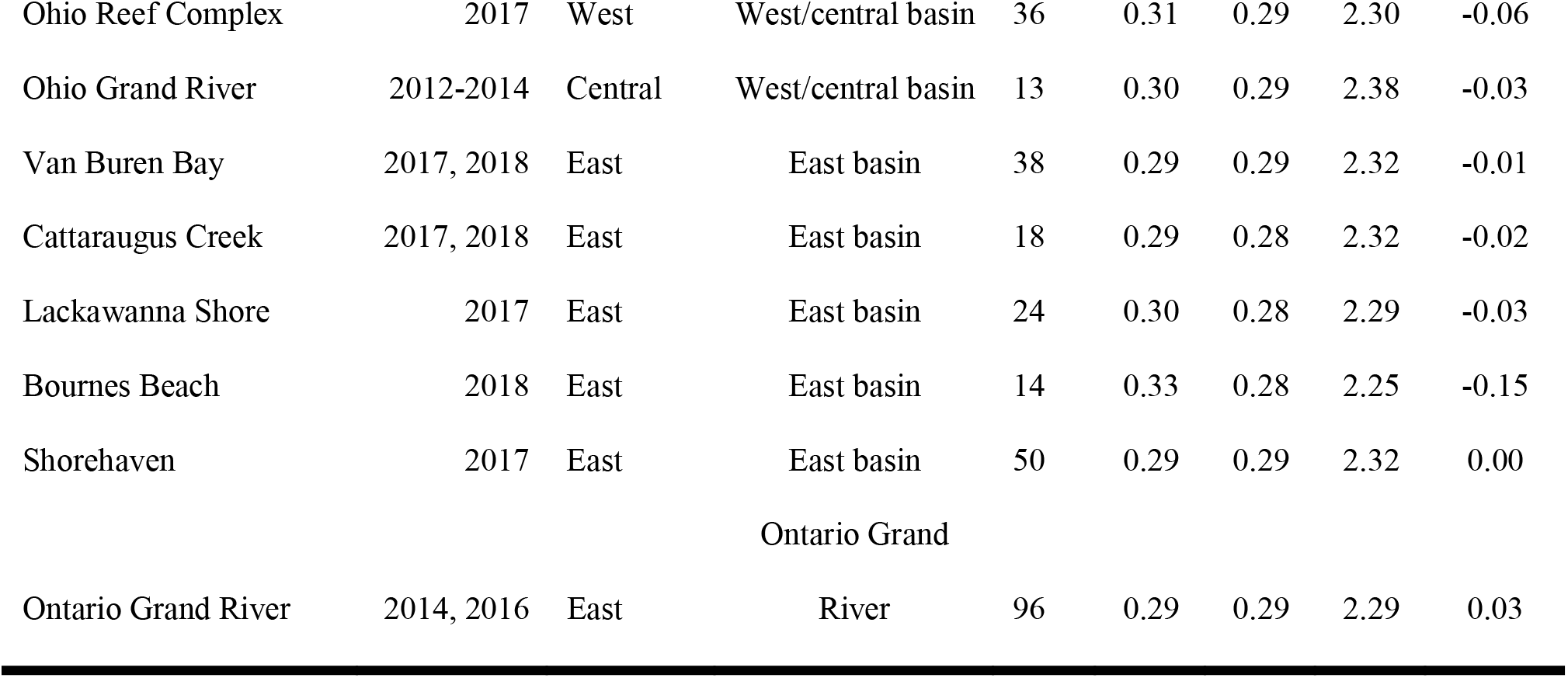
Walleye samples collected and genotyped for Rapture panel development and identification of reporting groups in mixed-stock analysis of recreational and commercial harvest in the East Basin of Lake Erie during 2016-2018. Columns show total sample size (N) and genetic diversity estimates including observed (H_O_) and expected (H_E_) heterozygosity, allelic richness (AR), and inbreeding coefficient (F_IS_) for final Rapture panel single nucleotide polymorphism (SNP) genotypes. These SNPs were identified during initial RAD-sequencing for panel development and microhaplotype genotypes identified through Rapture sequencing of 397 baseline individuals used for reporting group identification (note the higher AR of microhaplotype loci than SNP loci in Baseline individuals). *Panel Development* samples generated the initial RAD-sequencing data that were subsequently used to develop the Rapture panel. This panel, in turn, was used to describe the *Baseline* population structure in Lake Erie.

A single RAD-seq library was prepared using *SbfI* enzyme and the standard library preparation approach outlined in Ali et al. (2016) and detailed in Ackiss et al. (2020). Samples were sequenced using paired-end 150 BP sequencing on an Illumina HiSeq4000 at NovoGene (Sacramento, CA). Loci were then assembled *de novo* in STACKS v. 2.0 (Catchen et al. 2011; 2013) and a catalog of all putative loci was created in *cstacks* from data of all 96 individuals. Samples were demultiplexed using *process_radtags* (-e sbfI -i gzfastq -c -q -r --filter_illumina – bestrad), assembled *de novo* in *ustacks* (----disable-gapped, -m3, -M 3, -H, --max_locus_stacks 4, - -model_type bounded, --bound_high 0.05), matched in *sstacks* (----disable gapped), converted to bam files using *tsv2bam*, and genotyped in *gstacks*. Finally, genotypes were called for all SNPs with a minor allele count greater than three (----mac 3). We then removed putative paralogs identified in HDPlot (McKinney et al. 2017b), as well as loci with minor allele frequencies <0.01, heterozygosity <0.05, and genotype rate < 50%. Sequence data for the remaining loci that met our quality standards were sent to Arbor Biosciences (Ann Arbor, MI) for bait development. Arbor Biosciences conducted additional quality filters including a blast alignment to the yellow perch (*Perca flavescens*) genome (Feron et al. 2019) and then designed and synthesized two baits per-locus resulting in a final panel of 12,081 loci and 24,162 unique baits. STACKS 2 catalog files and a fasta file for all 12,081 baited loci can be found on Dryad (*data will be available upon acceptance*).

### Baseline sequencing

The baseline population structure among six east basin, four west basin, and one central basin walleye spawning stocks (Fig. 1) was evaluated by sequencing 397 individuals collected during the 2012-2018 spawning seasons using our Rapture panel described above (Table 1). All samples were collected and extracted under the same protocols as Rapture panel development. Whenever possible samples from at least two different years collected from the same site were used to account for possible temporal variation (Table 1). Individuals sampled on known spawning grounds during the peak spawning season were presumed to have spawned in those areas. RAD-seq libraries were prepared identically as panel development individuals and then bait capture was conducted for each library following the myBaits protocol v.4.01(https://arborbiosci.com/mybaits-manual/) with minor modifications. In short, RAD-seq libraries were hybridized with the bait mixture for 16 hours at 65°C and then amplified using 10 PCR cycles, universal primers, and an annealing temperature of 56°C. Final Rapture libraries were then purified with a 1X Ampure bead solution and submitted for sequencing on ½ of a S4 NovaSeq lane at NovoGene (Sacramento, CA). Data were processed using STACKS v.2.3 (Rochette et al. 2019) with identical parameters and catalog as panel development. Finally, loci were filtered using a locus-specific whitelist that included only the 12,081 loci included in the final Rapture panel. Microhaplotypes at each locus were identified in the *populations* step of STACKS and used for all downstream analysis. To ensure that SNPs used in microhaplotype genotypes were not the result of sequencing error, loci had to be genotyped in at least 80% of individuals (both baseline and mixed-stock) and SNPs had to have a minor allele count of three or more to be processed.

### Baseline population structure and identification of reporting groups (natal sources)

*F*-statistics and discriminant analysis of principal components (DAPC; Jombart et al. 2010) were used to describe overall patterns of genetic structure in Lake Erie and to identify putative reporting groups for reassignment tests to determine classification accuracies. Observed and expected heterozygosity, inbreeding coefficient, and pairwise Weir and Cockerham *F*_ST_ were estimated in the DiveRsity v.1.1.9 R package (Keenan et al. 2013; R Core Team 2019). We used DAPC to identify putative clusters of spawning sites that could be combined into reporting groups for assignment of individuals of unknown origin. DAPC was run using the ADEGENET v.2.1.2 R package first with individuals grouped by spawning stock and next with individuals grouped into four clusters identified with the find.clusters() function (Jombart 2008). To avoid model over-fitting, the optimal number of principal components necessary to explain the variance among groups was identified in ADEGENET with the *optim.a.score*() function (Jombart 2008). Finally, we used AMOVA and permutation tests of significance implemented in the poppr v.2.8.3 R package to determine how much variance was explained when sites were grouped by putative reporting groups (Kamvar et al. 2014).

We tested reassignment accuracy of putative reporting groups identified through baseline population genetic analysis in the assignPOP v.1.1.8 R package (Chen et al. 2018). The assignment tests were performed using Monte-Carlo cross-validation in which individuals were randomly resampled as a training set, using remaining individuals as a test (holdout) set to avoid upwardly biased test results (Anderson 2010, Waples 2010). Specifically, we chose three proportions (0.5, 0.7, and 0.9) of individuals from each reporting group and used either random half or all loci as training data (combination N = 6) to perform the assignment test. Each combination of training data and the test were iterated 30 times for a total of 180 assignment tests. This procedure allowed us to evaluate variation in assignment accuracy and how different proportions of training individuals influenced the assignment results.

The assignPOP predictive models were built using a support vector machine (with linear kernel and parameter cost = 1), which has been shown to generate higher assignment accuracies than other models (i.e., LDA, naiveBayes, decision tree, and random forest; Chen et al. 2018) and had the highest accuracy in preliminary trial runs. We ran the reassignment test twice: once with every spawning stock kept separate and once with stocks clustered into our reporting groups identified with DAPC and pairwise *F_ST_*. Because individual assignments can sometimes bias mixture results, we compared assignment accuracy identified in assignPOP with 100% mixture assignments tests conducted in the rubias R package v.0.3.0 (Moran and Anderson 2019). Assignment accuracy of 100% mixtures for all reporting groups were estimated using a leave-one-out approach, 25 replicates, and a mixture size of 100. All presented graphics were constructed in R primarily using the ggplot 3.3.0 (Wickham 2009), ggpubr v.0.2.5 (Kassambara 2020), ggsci v.2.9 (Xiao 2018), and the sf v.0.7-7 (Pebesma 2018), and scatterpie v.0.1.4 (Yu 2019) packages.

### Mixed-stock assignment of harvested walleye

To determine the relative contribution of Lake Erie’s walleye stock reporting groups to the east basin commercial and recreational harvest, 1,274 individuals of unknown origin were sampled during the spring, summer, and fall of the 2016-2018 fishing seasons (Table 2). For both commercial and recreational fishery samples, anglers and fishers reported the grid location of their harvest on a map provided by creel agents at boat ramps/docks. The total length (nearest 1 mm) of all sampled individuals was also recorded. To maximize our ability to estimate seasonal variance in stock structure, the majority of our mixed-stock samples (N=1,021, 80%) were taken from the 2017 harvest that occurred between May and December (Table 2). However, 130 samples were selected from the July 2016 commercial harvest and 122 from the July 2018 recreational harvest to investigate interannual variation. Fin clips were preserved in 95% ethanol until DNA extraction using Qiagen DNEasy 96 kits.

**Table 2:**
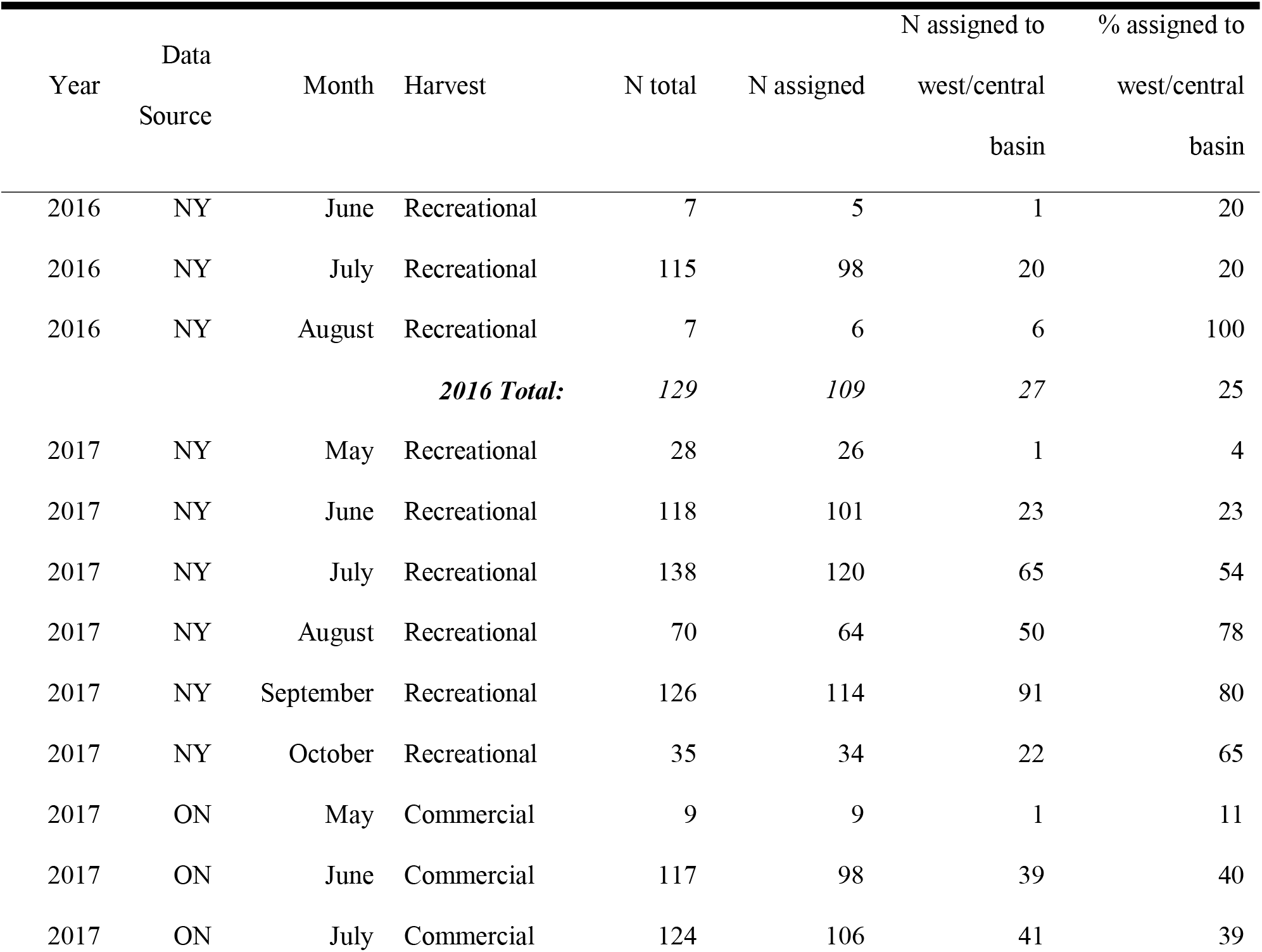
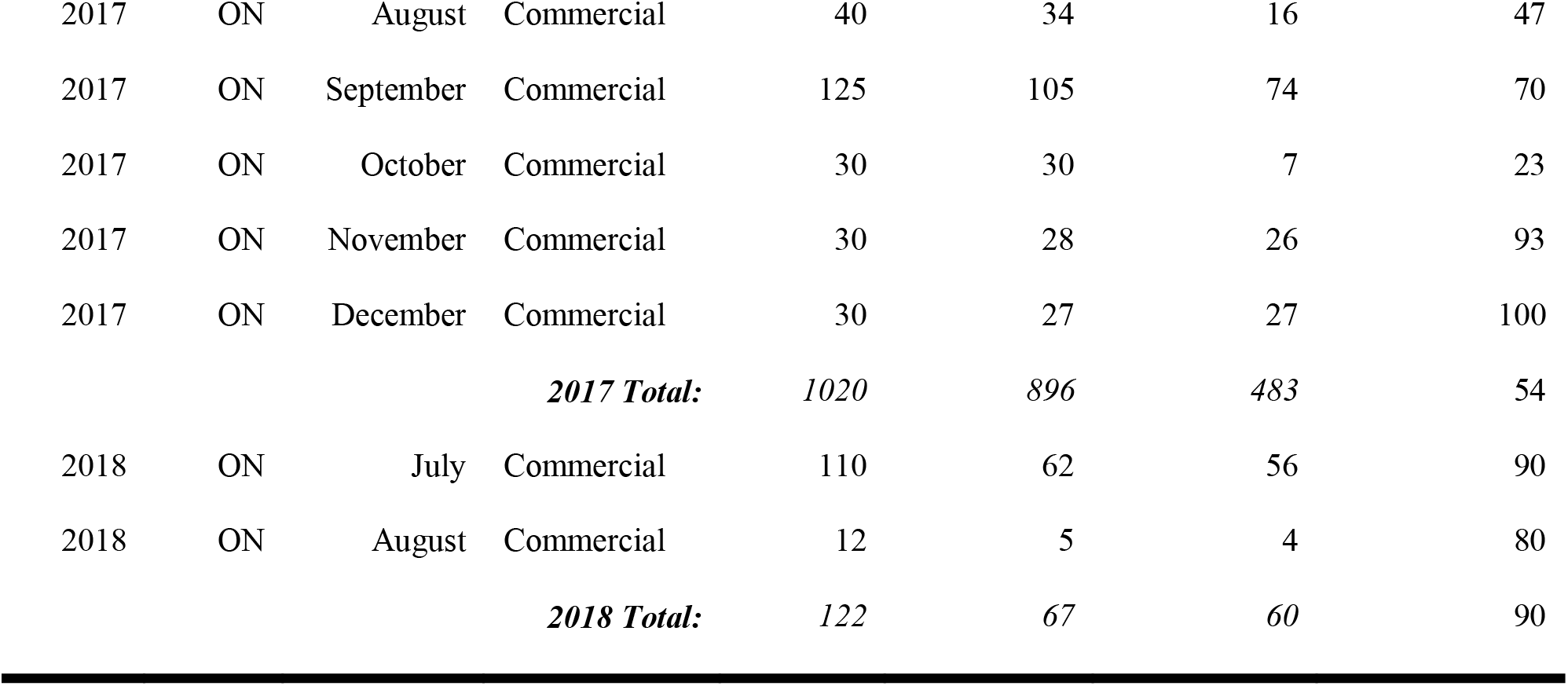
Number of walleye of unknown origin collected in the east basin of Lake Erie’s commercial and recreational fisheries that were genotyped (N total), successfully assigned to any one of three reporting groups: east basin, west/central basin, and Ontario Grand River (N reassigned), or assigned to West Basin (N assigned to west/central basin). Assignments to reporting groups were grouped by year collected, sample source (either New York Department of Environment and Conservation [NY] or the Ontario Ministry of Natural Resources [ON]), month of harvest and fishery type. Individuals that could not be assigned to a natal source had > 50% missing genotypes and were removed from analysis. For easier interpretation, the number of walleye reassigned to west/central basin is also reported as a percent of total fish assigned.

| Year | Data<br>Source | Month | Harvest | N total | N assigned | N assigned to | % assigned to |
| --- | --- | --- | --- | --- | --- | --- | --- |
|  |  |  |  |  |  | west/central<br>basin | west/central<br>basin |
| 2016 | NY | June | Recreational | 7 | 5 | 1 | 20 |
| 2016 | NY | July | Recreational | 115 | 98 | 20 | 20 |
| 2016 | NY | August | Recreational | 7 | 6 | 6 | 100 |
| **2016 Total:** |  |  |  | **129** | **109** | **27** | **25** |
| 2017 | NY | May | Recreational | 28 | 26 | 1 | 4 |
| 2017 | NY | June | Recreational | 118 | 101 | 23 | 23 |
| 2017 | NY | July | Recreational | 138 | 120 | 65 | 54 |
| 2017 | NY | August | Recreational | 70 | 64 | 50 | 78 |
| 2017 | NY | September | Recreational | 126 | 114 | 91 | 80 |
| 2017 | NY | October | Recreational | 35 | 34 | 22 | 65 |
| 2017 | ON | May | Commercial | 9 | 9 | 1 | 11 |
| 2017 | ON | June | Commercial | 117 | 98 | 39 | 40 |
| 2017 | ON | July | Commercial | 124 | 106 | 41 | 39 |
| 2017 | ON | August | Commercial | 40 | 34 | 16 | 47 |
| 2017 | ON | September | Commercial | 125 | 105 | 74 | 70 |
| 2017 | ON | October | Commercial | 30 | 30 | 7 | 23 |
| 2017 | ON | November | Commercial | 30 | 28 | 26 | 93 |
| 2017 | ON | December | Commercial | 30 | 27 | 27 | 100 |
| **2017 Total:** |  |  |  | *1020* | *896* | *483* | *54* |
| 2018 | ON | July | Commercial | 110 | 62 | 56 | 90 |
| 2018 | ON | August | Commercial | 12 | 5 | 4 | 80 |
| **2018 Total:** |  |  |  | *122* | *67* | *60* | *90* |

Rapture libraries for the mixed-stock analysis were constructed using identical procedures as baseline samples with one exception; to reduce the number of bait capture reactions necessary to genotype mixed-stock samples, DNA from two RAD-seq libraries were included in each bait capture reaction. All mixed-stock Rapture libraries were pooled and sequenced on four Illumina HighSeq4000 lanes at NovoGene (Sacramento, CA). Sequence data were processed in STACKS v.2.3 using identical procedures and filters as baseline samples with the exception of individual genotype rate, which was reduced to 50% because assignPOP indicated similar reassignment accuracy between 100% and 50% retained loci tests. Once all individuals were genotyped, each was assigned to one of the three reporting groups identified during baseline analysis using the assign.x function in assignPOP using microhaplotype genotypes and the SVM parameter. As an additional estimate of mixed-stock proportion in 2017, the total stock mixture was estimated in rubias with identical reporting groups and reference as assignPOP and bootstrapped proportions (100 bootstraps). As confirmation of individual assignments, we also used rubias to estimate the stock mixture of all individuals assigned to each reporting group. Individual assignments were presumed to be correct as long as the genetic mixture of individuals in each reporting group exceeded 90%.

### Stock-specific harvest dynamics

To determine how contributions to the eastern basin fisheries varied through space and time, all walleye that could be successfully assigned to one of our three reporting groups were summarized by year, month, location (commercial fishery or creel-grid centroid), and fishery type (recreational or commercial). For walleye sampled during 2017, the proportion of individuals assigned to each reporting group was calculated by dividing the number of walleye assigned to a particular reporting group by the total walleye assigned to any reporting group for a particular month, location, or fishery type. Because some locations contained a small number of assigned individuals in certain months, only month/location samples with greater than six successfully assigned walleye were included in these analyses (N = 48 grid by month samples). Using the proportion of walleye assigned to the west/central basin reporting group as our response variable, we constructed linear models using only 2017 data with fishery type, month, and longitude as predictor variables. To better identify patterns in harvest, longitude was used in place of grid identification numbers to orient positions along the east-west axis of Lake Erie. Estimates of stock proportion were also extrapolated to actual estimates of harvest (number of fish) for each fishery type, location, and month for 2017 (Walleye Task Group 2018). To describe inter-annual variation from 2016 to 2018, the proportional contributions of each reporting group during July was compared. Only samples collected during July 2016-2018 were used because July is consistently one of the peak months of walleye harvest in Lake Erie and we had a high number of available samples for all three years. These data were compared by calculating the proportion of individuals assigned to each reporting group relative to the total fish assigned to any reporting group that year. Because only a single month of samples was genotyped during 2016 and 2018, comparisons should be only considered as point estimates of proportional harvest.

Because the relative proportions of fish assigned to a particular reporting group throughout the season is most likely predicted by more than a single variable, we evaluated a series of seven generalized linear models (GLM) explaining the proportional harvest of walleye reporting groups. Information theoretic criteria (AIC_c_; Burnham and Anderson 2004) was used to identify a best model using available metadata for each fish: the type of fishery (commercial or recreational), average length of individuals sampled, latitude of capture, longitude of capture, and month of capture. All explanatory variables were first run independently against the response variable (proportion of walleye assigned to the west/central basin reporting group) and any variable not found to be significant (alpha = 0.05) was removed from future models. Once a subset of variables that were all independently significant were identified, all possible combinations of these variables were fit, and the best model was chosen based on the approach outlined in Symonds and Moussalli (2011). In short, each model was fit in R using the glm() function using default settings (Gaussian error structure) and then ranked based on AIC_c_ using the AICcmodavg v.2.3 R package whereby the model with the lowest AIC_c_ was considered to be the best fit. Finally, the Evidence Ratio (ER) was calculated in the qpcR v.1.4-1 R package as a measure of how much more likely the best model was than other models.

## Results

### Panel development

Panel development sequencing produced a total of 61,350,730 retained reads and an average effective per-sample coverage of 11.4 (standard deviation = 4.0). Of the 263,723 putative SNPs and 43,884 loci identified in the 96 individuals sequenced for panel development, 14,418 loci passed our quality control filters and were sent to Arbor Biosciences for bait development. Of the 14,418 sequences sent, baits were successfully designed for 12,081.

### Baseline stock structure

Using 395 walleye collected from spawning sites during the spawning season, we described the population structure of walleye spawning stocks in Lake Erie and identified three putative reporting groups. Baseline Rapture sequencing used to identify reporting groups produced a total of 1,953,890,992 reads and an average effective per-sample coverage of 124.2 (standard deviation = 44.2). Of the 12,091 baited loci, 8,482 loci passed the genotyping rate (80%) and minor allele count filters (minor allele count > 3). Microhaplotypes at each locus had an average of 3.8 alleles, ranging from 1 to 21 alleles. Two samples contained genotypes at fewer than 70% of loci and were removed prior to panel testing for a final total number of samples of 395 individuals and overall genotyping rate of 92%.

Three to four clusters of walleye spawning stocks that had similar allele frequencies were identified based on both pairwise *F*_ST_ (Table 3) and DAPC analyses (Fig. 1; Fig. 2). DAPC identified four distinct genetic clusters (Fig. 1, Fig. 2B). The first cluster identified by DAPC contained 142 out of 143 of the western basin and 13 out of 13 central basin walleye. The second cluster identified by DAPC contained 139 out of 142 individuals collected from 5 of the 6 eastern basin stocks sampled. Finally, individuals from Ontario’s Grand River in the eastern basin formed the remaining two clusters with 24 individuals assigned to cluster 3, 56 individuals assigned to cluster 4. Additionally, 16 individuals were genetically similar to west/central basin individuals and assigned to cluster 1 (Fig. 2). This same pattern was apparent with pairwise *F*_ST_ values (Table 3) whereby pairwise *F*_ST_ was highest between Ontario’s Grand River and all other sites in the eastern basin and all of the western basin sites (average *F*_ST_ = 0.032). Pairwise *F*_ST_ was lower for all non-Ontario Grand River comparisons but higher between basins (mean pairwise *F*_ST_ = 0.007) than within them (mean pairwise *F*_ST_ = 0.003).

**Table 3:**
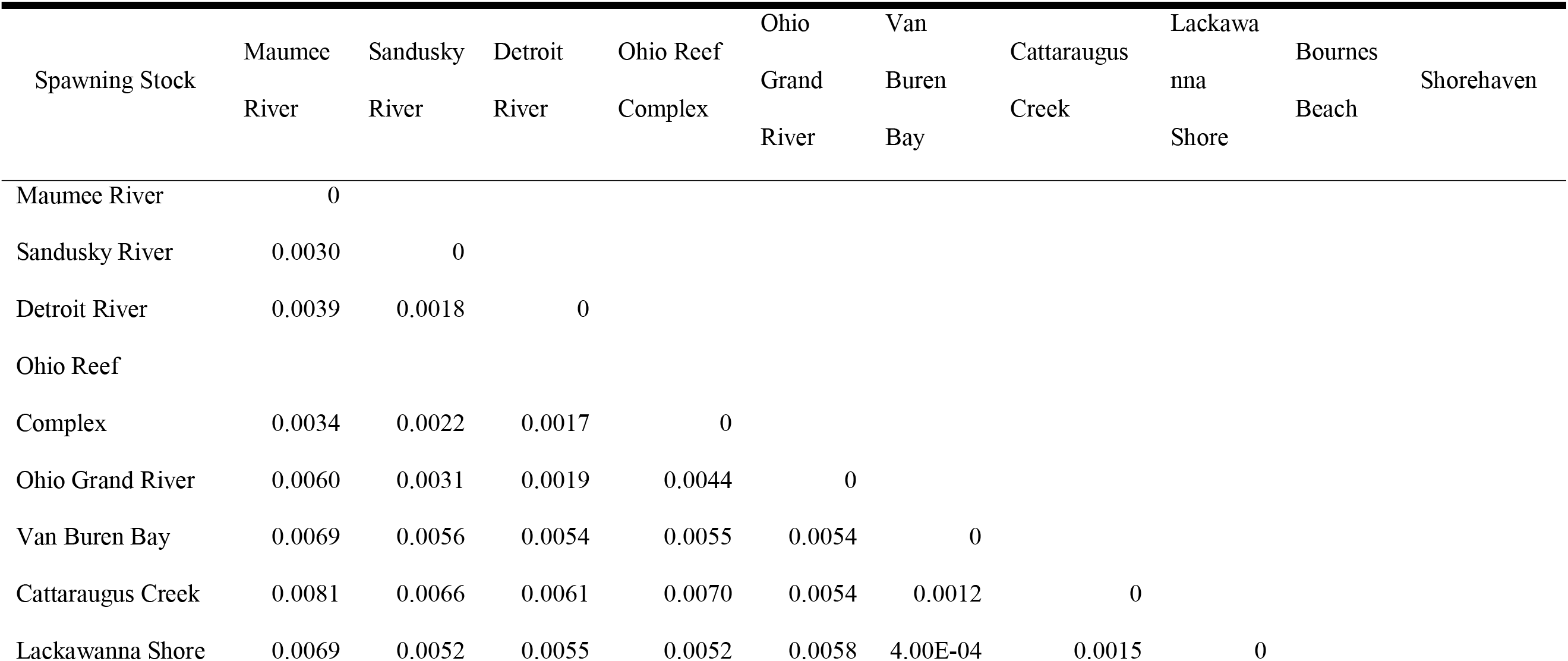
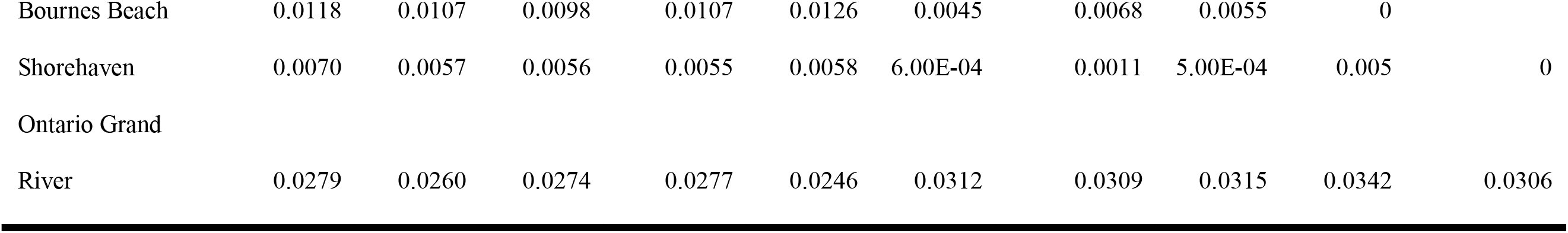
Pairwise genetic differences of walleye collected at different Lake Erie spawning locations during 2014 − 2017 (*F*_ST_; Weir and Cockerham, 1984). Individuals from all spawning stocks were used in the baseline assessment of population structure of Lake Erie and used to define reporting groups for mixed-stock analysis.

**Fig 2:**
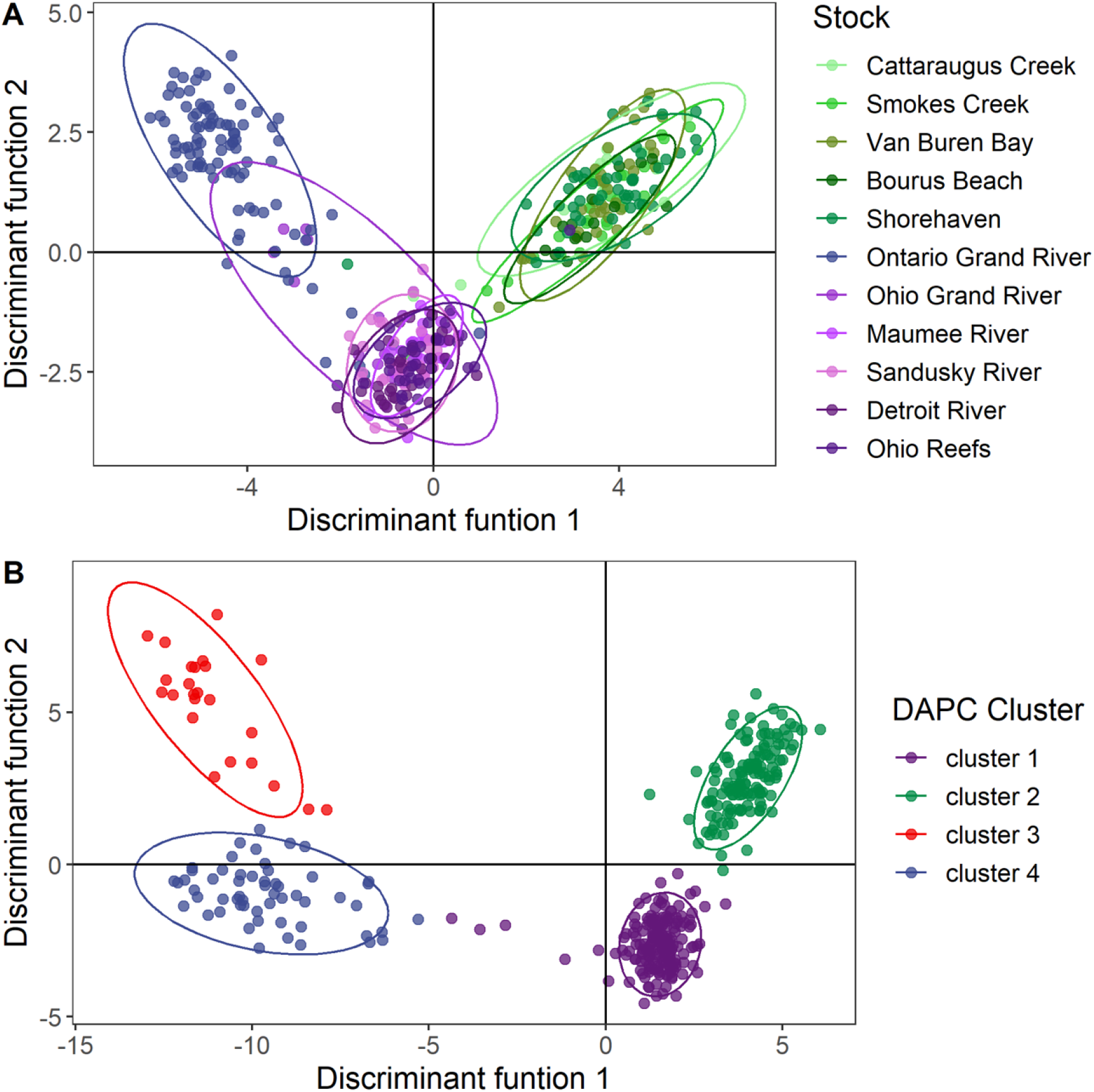
Discriminant analysis of principal components (DAPC) of walleye (individual points) collected on spawning grounds during the spawning season 2012 –2018. Individuals are grouped by spawning stocks (A) and into the 4 most parsimonious clusters based on Bayesian Information Criterion (B). Ellipses show the 95% confidence interval around each group. Colors in A correspond approximately to the reporting groups shown in Figure 1: west/central basin = purples; east basin = greens; Ontario Grand River = blue.

Reassignment accuracy to individual spawning stocks was low (< 50 – 75% for most stocks; Suppl. Fig 1). Thus, based on DAPC and pairwise *F*_ST_, we decided to group spawning stocks into three reporting groups hereafter referred to as the west/central basin (containing all individuals from western basin and central basin stocks), east basin (containing individuals from 5 out of 6 eastern basin stocks), and Ontario’s Grand River (containing individuals from the Ontario Grand River stock). When spawning stocks were classified into these three reporting groups, a significant amount of variance was explained (AMOVA p = 0.01; variance between reporting groups = 1.9%, variance between populations within reporting group = 0.2%). The average overall reassignment accuracy to the west/central basin, east basin, and Ontario Grand River reporting groups was estimated to be 93% (Fig. 3). Reassignment to each reporting group varied and was lowest for Ontario Grand River (mean = 85%), and more similar between the east basin and west/central basin reporting group (mean = 96% and mean = 99% respectively). Reassignment accuracy was similar for all sets of training individuals and when either 50% or 100% of loci were used (Fig. 3). Baseline assessment with rubias found similar reporting group accuracy as assignPOP based on posterior mean reporting group scores (Suppl. Fig. 2). Thus, we have high confidence in using our three reporting groups to identify the source origins of walleye harvested in eastern Lake Erie’s fisheries.

**Fig 3:**
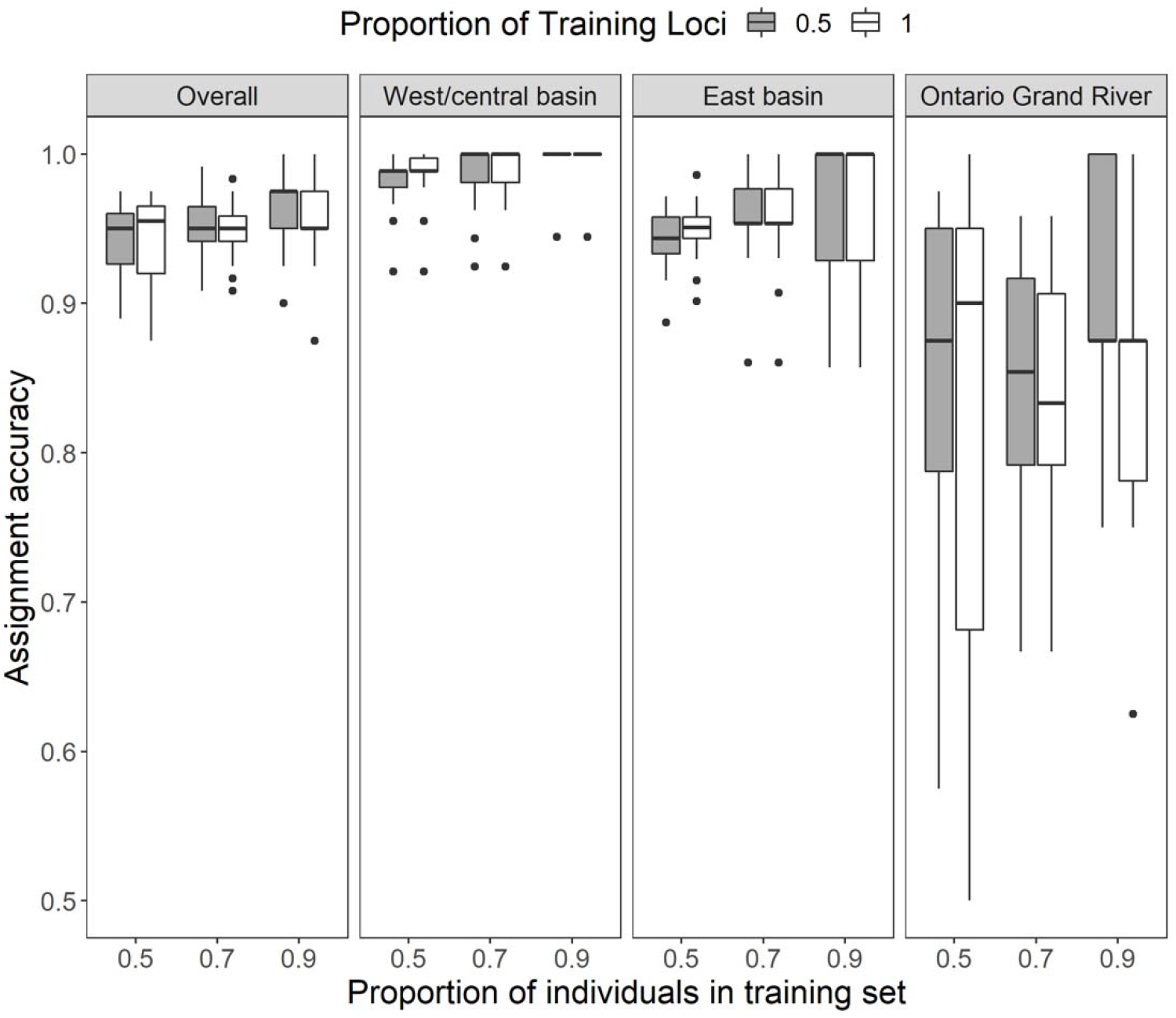
Reassignment accuracies of three reporting groups identified based on 395 adult Walleye collected from 11 Lake Erie spawning sites during the spawning season between 2012-2017. The reporting groups are (1) West/Central Basin: Maumee River, Sandusky River, Detroit River, the Ohio reef complex, Ohio Grand River; (2) East Basin: Shorehaven, Bournes Beach, Van Buren Bay, Cattaraugus Creek, Lackawanna Shore; and (3) Ontario Grand River. Reassignment accuracy was determined using either 0.5 or 1 proportion of training loci (gray and white bars, respectively) and a support-vector machine algorithm (Chen et al. 2020), with training samples for each grouping consisting of 0.5, 0.7, or 0.9 proportion of the collected individuals (chosen randomly). The remainder of individuals (0.5, 0.3, or 0.1) was used as the test (holdout) data set to determine reassignment accuracy. Box plots portray medians (thick black line), interquartile ranges (ends of boxes), and outliers (black dots).

### Assignment of mixed-stock individuals of unknown origin

Rapture sequencing of the mixed-stock walleye samples of unknown origin produced a total of 3,331,974,311 retained reads and an average effective per-sample coverage of 41.8 (standard deviation = 24.3). Of the 12,081 baited loci, 8,482 loci passed genotyping rate and minor allele count filters and overlapped with loci used for baseline analysis. Of the 1,274 individuals analyzed, 199 (15%) failed to genotype in at least 50% of loci and were removed from analysis (Table 2). However, because the removed samples were spread across sampling dates and locations, we do not believe that their removal led to important sampling bias or was the result of consistent laboratory error. All individuals assigned to either west/central basin or east basin reporting groups and no individuals assigned to the Ontario Grand River reporting group. During 2017, the percentage of individuals of west/central basin origin was identical between individual assignments determined in assignPOP and stock mixtures estimated in rubias (west/central basin = 54%; east basin = 46%; Ontario Grand River = 0%). Owing to the variable sample sizes among collections and similarity between assignPOP and rubias, we limited our subsequent analysis to individual assignments. The percentage of west/central basin origin walleye was almost identical between recreational and commercial fisheries (51% and 49%, respectively) indicating that both fisheries were equally supported by fish originating in west/central and east basin reporting groups (F = 0.1; P = 0.71; Fig. 4A).

**Fig 4:**
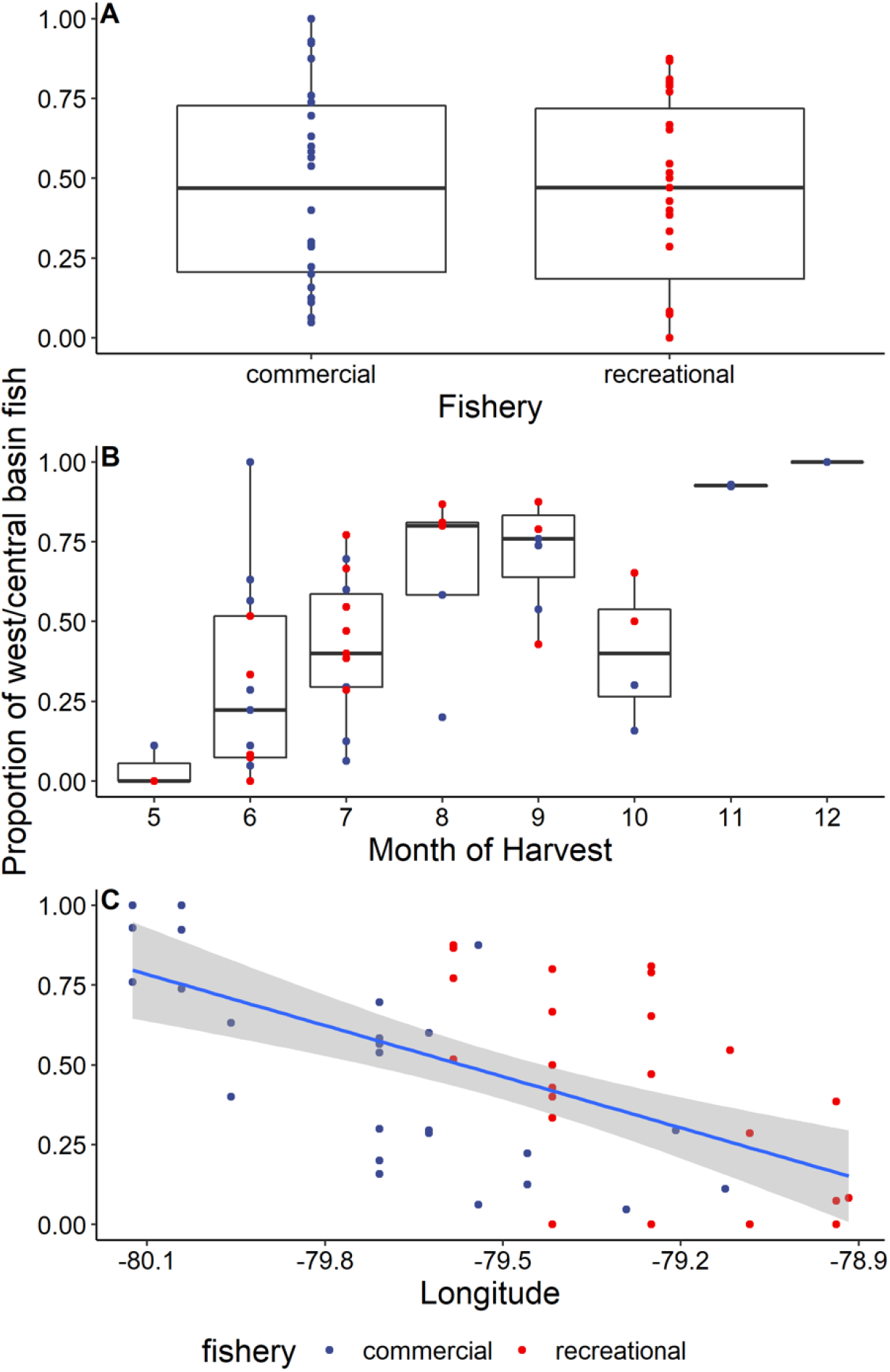
Proportion of walleye harvested in eastern Lake Erie’s recreational and commercial fisheries during 2017 assigned to the west/central basin reporting group (A), broken down by harvest month (B) and location (longitude; C). Each colored point represents the proportion of fish assigned to the west/central basin from a particular sampling event (grid/date combination). Blue line and gray background in C represent correlation of generalized linear model and 95% confidence interval. Only sampling events that contained greater than 6 assigned individuals were included. Note higher sample sizes for core months of harvest, June –September.

Although about half of all fish harvested in the east basin were found to be of west/central basin origin, the percent of west/central basin origin walleye harvested varied spatially and temporally (Fig. 4B; Fig. 5). During the spring, the proportion of west/central basin origin walleye in the harvest was low with increasing contributions from this reporting group throughout the summer (F= 25.1; P < 0.01). For example, 6% of walleye were of west/central basin origin in May 2017, which increased to 31% by the end of June 2017 (Fig. 4B). The presence of walleye of west/central basin origin in the commercial fishery remained high throughout the summer and into the fall (July – October average percent of west/central basin origin = 57%) Harvest composition also varied spatially, with more easterly sites having fewer individuals of west/central basin origin than more westwardly ones (F= 24.0; P < 0.01; Fig. 4C).

**Fig 5:**
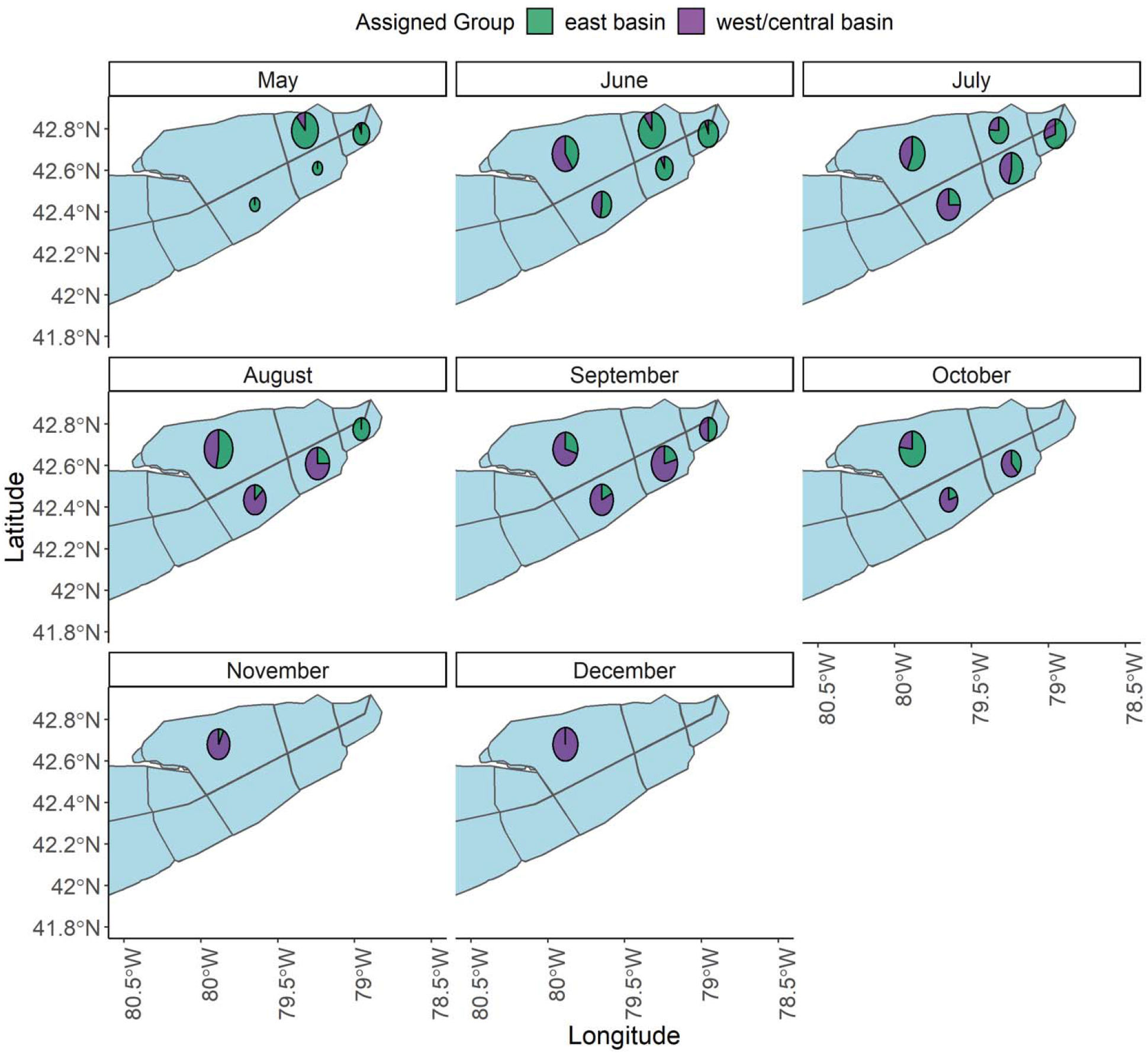
Monthly assignment of walleye to basin of origin sampled from the commercial and recreational harvest in Lake Erie’s east basin in 2017 from eight harvest zones. The size of each pie corresponds to log_10_ normalized mean number of fish harvested from creel sampling locations with centroids located within each harvest zone. Only two colors are shown because no harvested individuals were assigned to the Ontario Grand River reporting group.

Based on examination of samples collected during July 2016-2018 harvest composition varied among years (Table 2). The percentage of west/central basin fish in the July 2016 recreational fishery was 20%, which was smaller than the percentage of west/central basin origin fish harvested in the recreational (54%) or commercial (38%) fisheries during 2017. However, during July 2018 the percentage of walleye of west/central basin origin in the commercial harvest was 90%, which was higher than either the commercial or recreational harvest during July the previous years.

### Predicting contributions from the west/central basins

Modeling showed that the proportional contribution of the western/central basin spawning stocks to the eastern basin fisheries in 2017 varied both seasonally and spatially. The candidate model with the highest level of support in our AIC_c_ contained three of the five potential predictors (longitude, latitude, and month), and had an evidence ratio that indicated it was 10 times more likely than the next best model (Table 4; AIC_c_ = −5.5; degrees freedom = 47; Intercept = 2.8; latitude = −0.74, longitude = −;0.37, date = 0.05). Fishery type and mean total length were not significant predictors of proportion of west/central basin fish, and so were removed prior to AIC_c_ analysis. All other variables appeared to explain a significant amount of the variance and were included in our AIC_c_ assessment. Longitude was found to be the most important predictor of proportion of west/central basin walleye with proportional contributions of west/central basin walleye in harvest declining from west to east (Fig. 4C), however date and latitude were also important based on ER and delta AIC_c_ with proportional contributions of west/central basin walleye increasing from spring to fall and decreasing from North to South.

**Table 4:** Statistics from comparisons of general linear models used to describe the proportion of west basin walleye (PropWB) harvested in the east basin of Lake Erie’s recreational and commercial fisheries during 2017. The number of estimated parameters for each model (K), Akaike’s information criterion for small sample sizes (AIC_c_), change in information criterion between sequential models (Delta AIC_c_), AIC_c_ weight (AIC_c_Wt), and evidence ratio (ER) are provided. The most parsimonious model is in bold-face text

| Model | K | $AIC_c$ | Delta $AIC_c$ | $AIC_cWt$ | ER |
| --- | --- | --- | --- | --- | --- |
| **PropWB~lon+lat+month** | **5** | **-5.54** | **-** | **0.869** | **-** |
| PropWB~lon+lat | 4 | -0.99 | 4.52 | 0.091 | 10.14 |
| PropWB~lon+month | 4 | 1.28 | 6.79 | 0.029 | 31.52 |
| PropWB~lat+month | 4 | 3.64 | 9.15 | 0.009 | 102.57 |
| PropWB~lon | 3 | 7.61 | 13.12 | 0.001 | 788.41 |
| PropWB~month | 3 | 8.33 | 13.84 | 0.001 | 1130.40 |
| PropWB~lat | 3 | 19.11 | 24.62 | 3.92E-06 | 247246.43 |

## Discussion

Our study provides the first successful attempt to use a sequencing-based genetic panel to determine the relative contributions of local spawning stocks to harvest in the Great Lakes. Walleye of unknown origin harvested by commercial and recreational fisheries in eastern Lake Erie were assigned to one of three reporting groups with high accuracy using a Rapture panel and genotype data from 8,482 microhaplotype markers. Results supported the expectation that walleye from the productive west/central basin spawning stocks contributed substantially to the eastern basin fisheries for much of the year. Even so, seasonal and spatial variance in the proportion of walleye originating from the west/central basin indicated that the smaller eastern basin stocks (with the exception of the Ontario Grand River) comprised a large portion of harvest during certain times, such as the spring, or in certain parts of the basin, such as more easterly parts of the basin. In addition to providing critical information on stock-specific harvest of walleye in the eastern basin of Lake Erie that can benefit fishery management, our study represents one of the first uses of Rapture data for mixed-stock assignment in fisheries (but see Carrier et al. 2020) and the first in a freshwater ecosystem.

### Population structure and reassignment accuracy

By using microhaplotype genotypes at 8,482 loci, we were able to accurately (> 95%) assign fish of unknown origin to one of three reporting groups of walleye spawning stocks in Lake Erie. Our high assignment accuracy is an improvement relative to previous attempts at differentiating Lake Erie stocks of walleye using microsatellite loci (Stepien et al. 2012; Brenden et al. 2015) or mitochondrial haplotypes (Stepien and Faber 1998; Gatt et al. 2004; Haponski et al. 2014). Unlike our approach, which aggregated spawning stocks with low genetic differentiation, these earlier studies treated each spawning stock independently (Strange and Stepien 2007; Stepien et al. 2012), and often included only a subset of known stocks within a given basin (Brenden et al. 2015; Chen et al. 2019). In turn, while microsatellite loci and mitochondrial haplotypes identified significant genetic differences between the west and east basins, the reassignment accuracy to individual stocks in these studies was generally low (<25%) and inconsistent between western basin versus eastern basin reporting groups (20 – 87%). When taken in context with these previous studies, our results indicate that walleye of unknown spawning stock origin can accurately be assigned to one of three reporting groups (i.e., western/central basin, eastern basin, and Ontario Grand River) but not to a single spawning stock within a reporting group. The high reassignment accuracy in our study can be attributed to the increased statistical power attained by genotyping thousands of genomic markers and by conducting a comprehensive baseline assessment of lake-wide spawning stocks. Both of these factors made it possible to distinguish among weakly structured spawning stocks and provide a realistic representation of the interannual movement and mixture of walleye stocks in the eastern basin of Lake Erie during our study.

### Stock contributions to eastern basin fisheries

Walleye from the west/central basin substantially contributed to the recreational and commercial harvest in the eastern basin, although their contribution varied among years, seasons, and locations. Walleye from the western basin are known to migrate seasonally towards the eastern basin after spawning (Zhao et al. 2011; Vandergoot and Brenden 2014; Matley et al. 2020), but the extent to which these migrations influenced the actual harvest of walleye was previously unknown. Because western basin walleye spawning stocks are highly productive (DuFour et al. 2015) and have been tracked migrating to the eastern basin (Zhao et al. 2011; Matley et al. 2020), we were not surprised that western basin walleye contribute to eastern basin harvests. Importantly, our first quantification of seasonal, annual, and spatial variation in the contributions of western basin migrants to eastern basin fisheries supports previous evidence that walleye stocks can be treated as a portfolio whereby both eastern and western basin spawning stocks contribute to lake-wide harvest opportunities (DuFour et al. 2015). Further, these results provide an important first step toward a lake-wide approach to assessment and management of Lake Erie walleye, which was identified as a key priority in the current walleye management plan (Kayle et al. 2015).

The proportion of walleye of west/central basin origin caught in the eastern basin was low during the spring fishing season and increased rapidly following spring spawning period in April. By early summer (June – July) west/central basin walleye already made up nearly 50% of the harvest in our 2017 samples, and by August, the majority of genotyped fish in both the recreational and commercial harvest were of west/central basin origin. This finding is consistent with recent findings for large-scale acoustic telemetry studies examining annual migration patterns of Lake Erie walleye (Matley et al. 2020). Specifically, walleye from the western basin began being detected at central and eastern basin sites in higher proportions in June and continued to be detected throughout the summer (Matley et al. 2020). These annual movements have been hypothesized to be associated with such as thermoregulation (Raby et al. 2018) and prey availability (Kershner et al. 1999). Therefore, interannual variation in water temperature could influence the timing of spawning and subsequent eastward migration of west basin walleye which in turn could drive harvest dynamics (Dippold et al 2020). Similarly, changes in abundance of preferred forage fishes (e.g., emerald shiner *Notropis atherinoides*, rainbow smelt *Osermerus mordax*) may also influence migratory tendencies of western basin walleye. During our study, the proportion of west/central basin walleye harvested in the eastern basin during July varied from 20% in 2016 to 90% in 2018 across both fisheries (a 4.5-fold change), indicating interannual variation in stock composition can be high. This variation may be in part driven by stock-specific behavior and productivity. For instance, western basin walleye stocks more strongly selected depths > 13 m than did their eastern basin counterparts, suggesting that there could be potential for differential exposure to commercial and recreational fisheries (Matley et al. 2020). Similarly, in years when eastward migration is either delayed or limited, eastern basin stocks may be exploited to a higher degree (Dippold et al. 2020). Exploitation of individual stocks may also vary among years owing to differences in stock productivity and recruitment success for eastern and western basin stocks. For example, no fish in our mixed-stock analysis were assigned to the Ontario Grand River. However, this stock has been suggested to substantially contribute to eastern basin harvest in other years (C. Wilson, *unpublished data*). Therefore, periodic (e.g., every 3-5 years) reassessments of stock contributions to the harvest are likely necessary to allow management agencies to characterize spatio-temporal variation in relative stock contributions.

Even when walleye of west/central basin origin were present in the eastern basin, their contribution to harvest was spatially variable, with higher proportions of these individuals being recovered in more westerly areas of the eastern basin relative to more easterly sites. This trend was especially clear during June and July, where west/central origin fish made up 50 – 75% of assigned individuals in our westerly samples, whereas this percentage was closer to 25% in easterly samples (Fig. 5). This finding is also in accordance with previous studies of walleye movement, where the number of western basin walleye detected decreased with distance from the western basin (Raby et al. 2018; Matley et al. 2020). Month and longitude of harvest alone described over 60% of the variance in stock proportions during 2017, suggesting that the contribution of western basin walleye to the eastern basin harvest is non-random and that these movements should be considered in management of walleye. Understanding the migration patterns and environmental predictors of stock structure have proven crucial to defining harvest quotas in both Pacific and Atlantic salmon populations (Shaklee et al. 1999; Vähä et al. 2011; Bradbury et al. 2016). Once a baseline of genetic connectivity and movement is established, assessments of mixed-stock composition can help identify interannual shifts in abundance and help managers anticipate changes in stock abundance. Our findings support previous work documenting movement of western basin walleye into the eastern basin (e.g., Zhao et al. 2011; Matley et al. 2020) and advanced understanding by providing a baseline estimate of harvest composition in the eastern basin.

Eastern basin walleye stocks clearly do have an influence on eastern basin fishing opportunities and contribute to the stability of the Lake Erie walleye population year-to-year. For example, because the presence of west/central basin walleye decreases from west to east (see Fig. 4), recreational walleye fishing in the eastern most portion of the eastern basin is likely supported primarily by walleye of east basin origin for much of the year. Additionally, an estimated 11,694 walleye were harvested by commercial fishers in the east basin during May 2017 at a time when greater than 90% of walleye that we sampled originated from the east basin. During June, the estimated catch rose to 76,579, 40% of which may have originated from the smaller, less productive east basin spawning stocks (Suppl. Fig. 3). These simple extrapolations illustrate the utility of mixed-stock analysis for providing important information that can be used to protect small and vulnerable stocks such as those found in the eastern basin of Lake Erie. Further, understanding how and when smaller stocks are exploited in high numbers makes targeted action to limit overharvest possible (Dann et al. 2013), thus offering a means to help maintain a diverse portfolio of stocks that we expect to be critical for sustaining fisheries production in the face of continued anthropogenic change.

No walleye originating from the Ontario Grand River reporting group were identified in either fishery. While this finding could be real, the possibility also exists that either we failed to collect Ontario Grand River walleye in our samples or that Ontario Grand River individuals were mis-assigned in our analysis. The estimated reassignment accuracy of Ontario Grand River walleye was the lowest of our three reporting groups but was still high (87%) relative to other genetic studies of walleye in Lake Erie (Johnson et al. 2004). The variability in our assignment accuracy of the Ontario Grand River reporting group can be attributed to the 16 baseline individuals sampled from the Ontario Grand River that were genetically similar to the west/central basin reporting group. Therefore, any fish sampled that was genetically similar to the majority of Ontario Grand River walleye used in the baseline would most likely have been successfully assigned to this source stock, suggesting that misassignment is not the cause of failed detection. However, some behavioral evidence exists that suggest that walleye from Ontario’s Grand River remain close to the northern shore of Lake Erie throughout the summer, and do not mix evenly with walleye from the west/central and east basin reporting groups (TM, *unpublished data;* Matley et al. 2020). Therefore, our sample collections from commercial fisheries may not have included Ontario Grand River walleye. Additional assignment studies focused on sampling the Ontario Grand River reporting group could help to determine why this particular stock is unique from the rest of the Lake Erie walleye stocks, and help to verify if indeed the Ontario Grand River spawning stock does not significantly support the east basin recreational and commercial walleye harvest.

### Implications for inland fisheries conservation and management

By developing a Rapture panel, we were able to genotype a large number of individuals (1,671) at a consistent set of markers, which provided a clearer description of walleye genetic stock structure than was previously possible. The increased diagnostic power of our Rapture panel also allowed us to achieve higher confidence in individual assignments of natal origin, which is crucial for obtaining unbiased estimates of harvest proportions. When assignment accuracy is low, using individual assignments rather than mixture analysis for mixed-stock analysis can bias mixture estimates (Manel et al. 2005). However, the mixture proportions estimated based on our individual assignments were very similar to those estimated with traditional mixture analysis. Therefore, the ability to use thousands of loci for population assignment allowed us to obtain a high resolution and specificity of individual assignment without losing the precision of mixture analysis. Previous studies have shown that individual assignments can provide similar stock composition estimates as mixture analysis, but genetic differentiation among populations in these studies was larger than our study (Potvin and Bernatchez 2001). Indeed, when genetic differentiation is low, having a high number of genetic markers becomes more important for assignment accuracy (Waples and Gaggiotti 2006; Larson et al. 2014; Benestan et al. 2015). Genotyping-by-sequencing methods, like Rapture, are a feasible way to obtain this level of diagnostic power in a non-model species and can provide reliable mixture estimates. Once stock differences are identified, markers from high-density marker panels can be filtered to identify smaller subsets of highly-informative markers that can be incorporated into even higher throughput genotyping-in-thousands (GT-seq) panels for long-term monitoring and annual mixed-stock assessments (Bootsma et al. 2020). For this reason, we are optimistic that our approach will offer a new means for agencies managing fisheries with weak population structure to discern local spawning stocks such that their relative contributions can be quantified.

Our study provides the first estimate of proportional harvest of eastern and western basin walleye stocks, and therefore has direct implications for the sustainable management of this ecologically and economically important population (Hatch et al. 1987; Kayle et al. 2015). The baseline assessment of spawning stocks that we used to identify reporting groups (putative source stocks) refined previous assessments of walleye connectivity among spawning sites and helped to describe a third reporting group, the Ontario Grand River population, which had previously been suggested to be diverged from other eastern basin stocks (MacDougall et al. 2007). Subsequent mixed-stock assessments in the eastern basin identified clear patterns in harvest, helping to illuminate where and when different stocks were being exploited that would have otherwise been unattainable. Our results indicate that similar approaches could be used successfully in other ecosystems to obtain a snapshot of stock structure and exploitation without intensive up-front research. Harvest from rivers and lakes has been increasing worldwide (Fluet-Chouinard et al. 2018) but a lack of basic knowledge exists about many of these fisheries precludes the determination of necessary regulations and development of sustainable management plans (Allan et al. 2005; Bower et al. 2020). Determinations of relative stock contributions, like the one we present here, have the potential to inform sustainable harvest in many of these fisheries by identifying over-versus under-exploited stock components (Beard et al. 2011). This knowledge could open up the opportunity to identify stock components of other populations being stabilized by the portfolio effect and predict the sustainability of these populations long-term.

Genomics is emerging as an influential tool for managing and conserving freshwater fishes globally (Tibihika et al. 2020; Biesack et al. 2020). In Africa and Asia, genomics has become an important tool to quantify diversity and identify critically endangered populations (Malinsky et al. 2018; Ackiss et al. 2019). In Europe, genomics has become central to domestication of native species for aquaculture (Toomey et al. 2020). In the Laurentian Great Lakes, researchers have made substantial advances in the last decade to incorporate molecular techniques into the management and conservation native communities including lake trout *Salvelinus namaycush* (Bernatchez et al. 2016), brook trout *Salvelinus fontinalis* (Elias et al. 2018), coregonines (Ackiss et al. 2020), and walleye (Chen et al. 2019). Much of this success can be attributed to increased access to molecular resources for non-model organisms and reduced sequencing costs that have made large-scale genomic assessment feasible for freshwater fisheries that often lack resources (Lynch et al. 2016). With the promise that genomic tools hold, we are hopeful that molecular studies of exploited freshwater populations can move beyond describing the extent of genetic diversity present and population structure in the system and begin being used to consistently monitor contemporary changes in population structure and microevolution in response to anthropogenic change.

Our study highlights the ability for genomic techniques to be applied directly to active fisheries and to quantify contributions of genetically distinct units to harvest, even those that are weakly differentiated. Without substantial prior genomic resources, we successfully completed all steps of a mixed-stock analysis that facilitated the quantification of stock contributions to fisheries harvest and identified predictors of stock composition of a large fishery in a single study. Genotyping-by-sequencing approaches allowed us to identify informative markers in a non-model organism while the construction of a Rapture panel enabled the consistent genotyping of thousands of SNPs across a large number of individuals. Our approach offers a pathway to obtain a snapshot of mixed-stock composition that has the potential to inform stock connectivity and temporal and spatial changes in harvest for data-poor fisheries with weak population structure where an understanding of stock structure and relative contributions to the fishery has not been feasible but is needed (Allan et al. 2005; Irvine et al. 2019). In this way, we are confident that the continued use of genomic approaches, like the one we demonstrated with Lake Erie walleye, will help untangle the spatio-temporal contributions of spawning stock portfolios to fishery harvest.

## Acknowledgements

We thank Michael Sovic for his initial work writing the grant and developing pilot data, Kristen Gruenthal, Tina Werner, and Kevin Smith for help processing tissue samples and conducting lab work, and Zac Driscoll and Zachary Feiner for help with statistical analysis. This work was funded by the Ohio Sea Grant College Program (#NA18OAR417100 to SAL) and research was supported by the Research Computing clusters at Old Dominion University. Any use of trade, product, or firm names is for descriptive purposes only and does not imply endorsement by the U.S. Government. There is no conflict of interest declared in this article.

## Data Archiving Statement

The data that support the findings of this study are openly available in Geome at https://geome-db.org/workbench/project-overview?projectId=190 and Dryad at https://doi.org/10.5061/dryad.4b8gthtb2. All raw sequence files will be uploaded and stored on the NCBI Short Read Archive upon manuscript acceptance.

## Supporting Information

Suppl. Table 1: Locus specific diversity estimates (Locus ID, number of alleles, effective number of alleles, observed heterozygosity, inbreeding coefficient (G_IS_), and genetic distance estimate (G_ST_) for the 8,482 microhaplotypes used for individual assignments.

Suppl. Table 2: Metadata collected for all mixed-stock individuals including assigned basin.

Suppl. Fig. 1: Reassignment accuracies of 11 spawning stocks identified based on 395 adult Walleye collected from 11 Lake Erie spawning sites between 2012-2017 calculated in assignPOP: West/Central Basin –Maumee River (MA), Sandusky River (SA), Detroit River (DE), the Ohio reef complex (RE), Ohio Grand River (OHG), Ontario Grand River–Ontario Grand River (ONG), East Basin –Shorehaven (SH), Bourus Beach (BB), Van Buren Bay (VB), Cattaraugus Creek (CC), Smokes Creek (SC). Reassignment accuracy was determined using either 0.5 or 1 proportion of training loci (colors) and a support-vector machine algorithm, with training samples for each grouping consisting of 0.5, 0.7, or 0.9 proportion of the collected individuals (chosen randomly). The remainder of individuals (0.5, 0.3, or 0.1) was used as the test (holdout) data set to determine reassignment accuracy. Box plots portray medians (thick black line), interquartile ranges (ends of boxes), and outliers (black dots).

Suppl. Fig. 2: Summary of rubias assignment accuracy of three Lake Erie reporting groups Ontario Grand River (ONG), west/central basin (West_Basin) and east basin (East_Basin). Mean posterior mixing proportions of 100% mixtures for each reporting group across the first 25 iterations (top) and distribution of individual posterior reporting group proportion for simulated collections from 100% mixtures of each reporting group.

Suppl. Fig. 3: Total walleye harvest by commercial and recreational fishing in 2017 across the 98 commercial harvest reporting grids and 13 recreational reporting grids located in the study area as reported in the annual Lake Erie Walleye Task Group report for 2017.

