## Supplemental Figures for "Mixed-stock analysis in the age of genomics: Rapture genotyping enables evaluation of stock-specific exploitation in a freshwater fish population with weak genetic structure"

Figure S1: Reassignment accuracies of 11 spawning stocks identified based on 395 adult Walleye collected from 11 Lake Erie spawning sites between 2012–2017 calculated in assignPOP: West/Central Basin – Maumee River (MA), Sandusky River (SA), Detroit River (DE), the Ohio reef complex (RE), Ohio Grand River (OHG), Ontario Grand River– Ontario Grand River (ONG), East Basin – Shorehaven (SH), Bourus Beach (BB), Van Buren Bay (VB), Cattaraugus Creek (CC), Smokes Creek (SC). Reassignment accuracy was determined using either 0.5 or 1 proportion of training loci (colors) and a support-vector machine algorithm, with training samples for each grouping consisting of 0.5, 0.7, or 0.9 proportion of the collected individuals (chosen randomly). The remainder of individuals (0.5, 0.3, or 0.1) was used as the test (holdout) data set to determine reassignment accuracy. Box plots portray medians (thick black line), interquartile ranges (ends of boxes), and outliers (black dots).

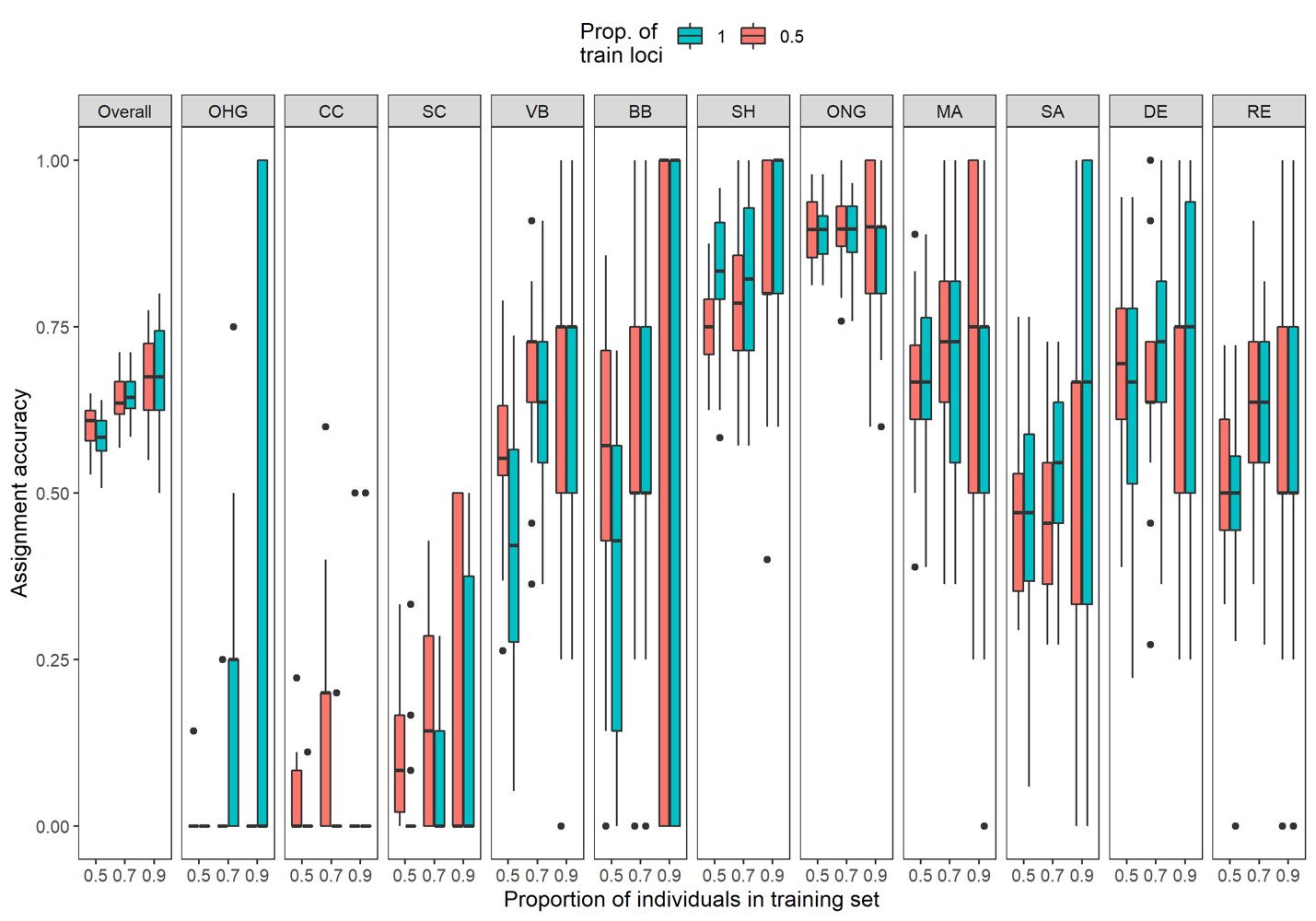

Figure S2: Summary of rubias assignment accuracy of three Lake Erie reporting groups Ontario Grand River (ONG), west/central basin (West_Basin) and east basin (East_Basin). Mean posterior mixing proportions of 100% mixtures for each reporting group across the first 25 iterations (top) and distribution of individual posterior reporting group proportion for simulated collections from 100% mixtures of each reporting group.

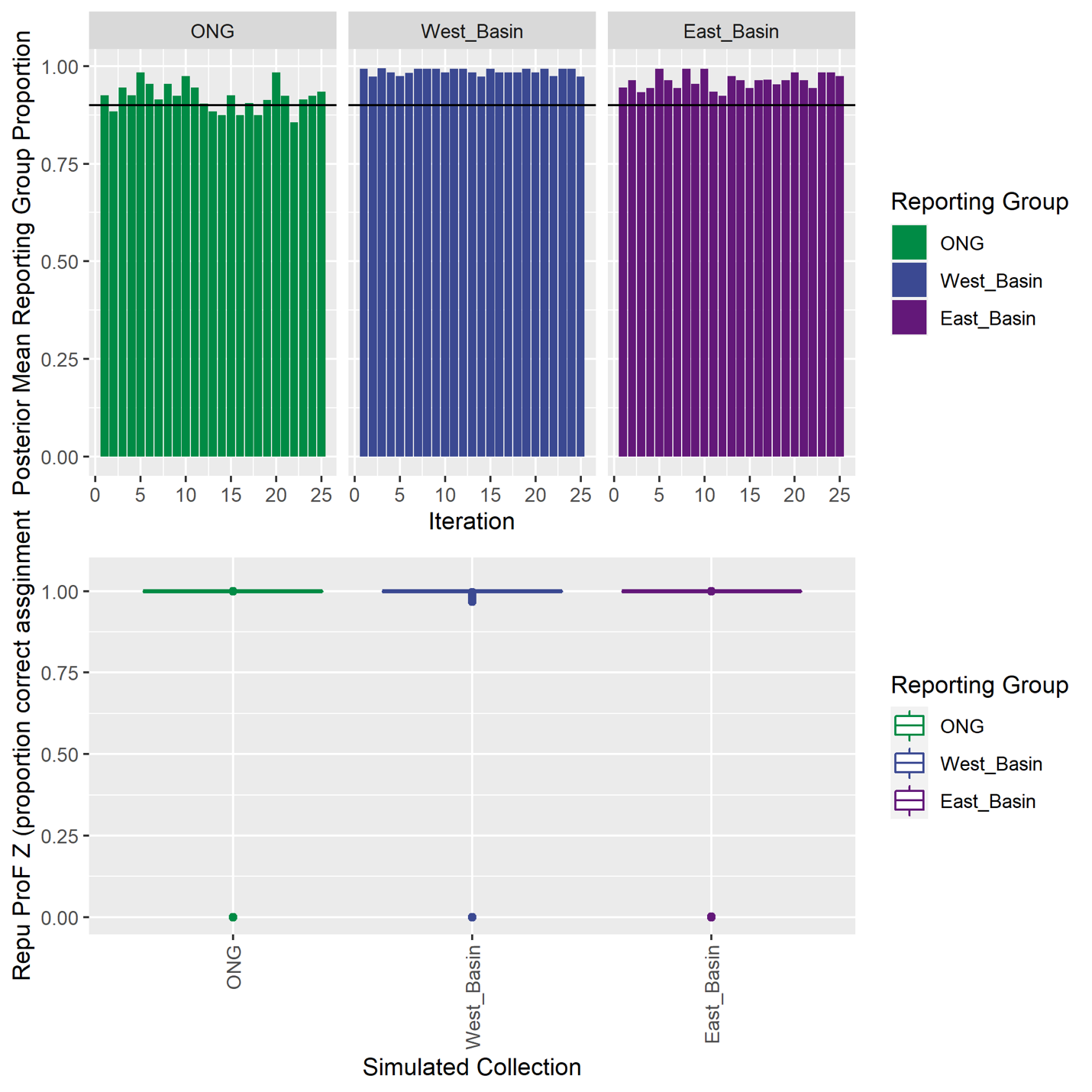

Figure S3: Total walleye harvest by commercial and recreational fishing in 2017 across the 98 commercial harvest reporting grids and 13 recreational reporting grids located in the study area as reported in the annual Lake Erie Walleye Task Group report for 2017.

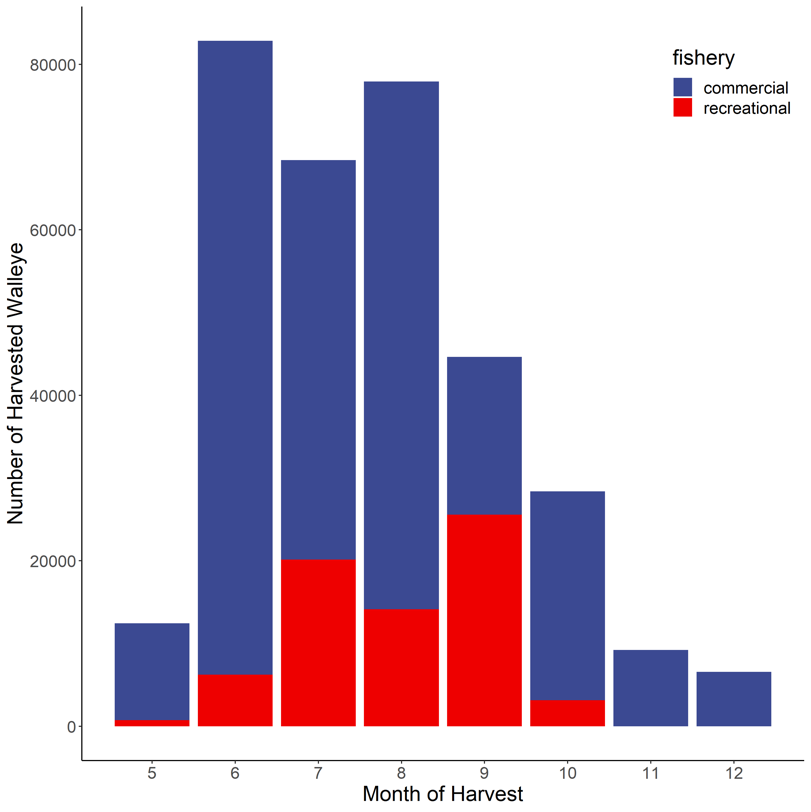
